# Repurposing Novobiocin for activity against latency associated *Mycobacterium tuberculosis* drug target nicotinate-nucleotide adenylyltransferase (Rv2421c)

**DOI:** 10.1101/2020.07.30.228726

**Authors:** Ruben Cloete, Mohd Shahbaaz, Melanie Grobbelaar, Samantha L. Sampson, Alan Christoffels

## Abstract

Nicotinamide-nucleotide adenylyl transferase (Rv2421c) was selected as a potential drug target, because it has been shown, *in vitro*, to be essential for *Mycobacterium tuberculosis* growth. It is conserved between mycobacterium species, is up-regulated during dormancy, has a known 3D crystal structure and has no known human homologs. A model of Rv2421c in complex with nicotinic acid adenine dinucleotide and magnesium ion was constructed and subject to virtual ligand screening against the Prestwick Chemical Library and the ZINC database, which yielded 155 potential hit molecules. 3D-QSAR studies of the 155 drug molecules indicated five compounds with similar inhibitory efficiencies compared to known inhibitors of Rv2421c. Molecular docking validation and molecular dynamics simulation analysis of the top five compounds indicated that the identified inhibitor molecules bind to Rv2421c with comparable efficiency as the substrate DND. Subsequent *in vitro* testing of the five compounds identified Novobiocin sodium salt with activity against *Mycobacterium tuberculosis* at 50 μM, 25μM and weakly at 10μM concentrations. Although, Novobiocin salt targets *Mycobacterium tuberculosis* DNA gyrase B our studies suggest that it has the potential to be repurposed to inhibit Rv2421c. Subsequent *in silico* structural analysis of known Novobiocin sodium salt derivatives against Rv2421c suggest promising alternatives for the treatment of *Mycobacterium tuberculosis*.

**Author Summary:** Rv2421c has been shown to be essential for *Mycobacterium tuberculosis* growth, shares no homology to known proteins in the human host, is conserved between various Mycobacterium species, is up-regulated during the non-replicative metabolic growth phase, making it an attractive drug target. It has a known 3D structure which has been exploited to screen for putative compounds within the Prestwick chemical library and ZINC database, resulting in the successful identification of 155 candidate compounds. Thereafter 3D-QSAR, molecular docking and molecular dynamics simulation studies were used to prioritize five potential compounds. Of the five compounds tested *in vitro*, only one, a Novobiocin disodium salt, showed activity against *Mycobacterium tuberculosis* at 50, 25 and weakly at 10 μM concentrations. Novobiocin is known to target *Mycobacterium tuberculosis* DNA gyrase B, but emerging resistance stimulated us to seek derivatives to target Rv2421c as alternatives for the treatment of *Mycobacterium tuberculosis*. Docking studies supported the higher binding affinities of Novobiocin derivatives to Rv2421c compared to DNA gyrase B. Future studies will involve testing these Novobiocin derivatives for activity against *Mycobacterium tuberculosis*.

## Introduction

Globally, Tuberculosis (TB) is the second leading cause of death after Human Immunodeficiency virus and accounts for approximately 1.3 million deaths and 10.4 million new cases per year [1, 2]. It is estimated that one-third of the world population is infected with TB. However, only about 10% of infected individuals develop active TB [3]. The remainder of latently infected individuals generate a critical reservoir of TB bacteria in their system which may result in disease reactivation should their immune system be compromised [3]. This poses a significant problem as current drugs require a long course of therapy with patients experiencing various degrees of adverse reactions [2]. Poor adherence due to adverse reactions is one of the main reasons patients default which may lead to further emergence of drug resistance. Furthermore, drug resistant strains of TB are on the rise warranting the urgent development of new drugs to combat this disease before it becomes an epidemic. The sequencing of several strains of *Mycobacterium tuberculosis* (*M. tuberculosis*) provides a useful resource to interrogate novel drug targets. The genome of *M. tuberculosis* is ∼4.4Mb in size and codes for approximately 4000 proteins [4]. However, not all these proteins are potential drug targets. For a candidate to be considered a promising drug target, it needs to meet certain criteria including being essential with regards to growth, replication and survival and not have any well-conserved homolog within its human host to avoid toxicity. Nicotinamide-nucleotide adenylyl transferase (NadD/Rv2421c) is essential for the survival of *Staphylococcus aureus, Streptococcus pneumoniae, Escherichia coli* (*E. coli*) and *M. tuberculosis* as all these organisms harbor both *denovo* and salvage pathways which is particularly useful during ATP limited conditions [5]. Furthermore, the NadD bacterial enzyme has a different 3D active site conformation compared to the human NadD isoform, providing an opportunity for selective inhibition. A previous study by Sorci and colleagues [5] used *in silico* screening and *in vitro* assays to identify structurally diverse compounds that inhibited NadD enzyme activity and cell growth in *E. coli* and *Bacillus anthracis* (*B. anthracis*). In a follow up study, Huang et al. [6] solved crystal structures of three inhibitors bound to NadD in *B. anthracis* that were identified in the Sorci study [5] and revealed a common binding site near residues Trp117, Tyr112 and Met109.

The enzyme NadD catalyses the transfer of an adenyl group from ATP to NaMN to form NaAD that is then converted to the ubiquitous intermediate nicotinamide adenine dinucleotide (NAD). Metabolic pathway and sequence similarity analyses indicated that Rv2421c is involved in nicotinate and nicotinamide metabolism in *M. tuberculosis* and has no equivalent human ortholog [7]. It is upregulated during non-growing metabolically active conditions of *M. tuberculosis* survival and lacks a close homolog in mice making it an attractive drug target [7]. The crystal structure of Rv2421c has been resolved and offers a unique opportunity for structure-based drug design [8]. Since, Rv2421c (PDBID: 4X0E) does not have ligands bound, we used the homologous structure of Nicotinic acid mononucleotide (NaMN) adenylyltransferase from *B. anthracis*, PDBID: 3E27 in complex with native substrate nicotinic acid adenine dinucleotide (DND) and magnesium ion (MG), to construct the structure of the Rv2421c in complex with DND and MG and then used this information to delineate the binding site of DND which was subsequently used for docking studies. We further used molecular dynamics (MD) simulation studies to validate binding free energy of the protein-ligand complexes to prioritize potential lead compounds. This was followed by experimental validation of the compounds as potential inhibitors of *M. tuberculosis* growth using *in vitro* growth assays.

## Materials and methods

### Structure preparation and assessment of Rv2421c (4X0E)

The crystal structure of Rv2421c (4X0E; resolution 2.4 Å, [8]) was superimposed onto the homologous template (the two proteins shared 40% sequence identity) to the structure of NaMN adenylyltransferase (PDBID: 3E27, resolution 2.2 Å, [5]) and the coordinates of substrate DND and MG were extracted from 3E27 to construct a complex of Rv2421c with DND and MG; the structural similarity between Rv2421c and 3E27 was assessed using the root mean square deviation (RMSD) value. Missing residues in the crystal structure of Rv2421c were modelled using the Swissmodel webserver [9]. The resulting complex of Rv2421c in complex with DND and MG was energy minimized using Desmond Schrodinger [10] for 2000 iterations of steepest descent method using the OPLS3 Force Field [11] in a TIP3 water box.

### Pharmacophore-based virtual screening

The energy minimized Rv2421c-DND-MG complex structure was used to generate a pharmacophore model using the program LIGANDSCOUT [13]. Two pharmacophore models were generated based on the substrate DND. The first model contained three features (H-bonds (carboxylate group), hydrophobic aromatic ring and negative terminal ionic phosphate group) which are conserved for DND interactions and also observed in the homologous template 3E27 (Fig 1A). The second model contained two features (three H-bonds (amine and hydroxyl group) and two hydrophobic aromatic ring contacts also important for DND interactions (Fig 1B). Subsequently, the generated pharmacophore models were screened against 1278 compounds from the Prestwick Chemical Library (PCL) (http://www.prestwickchemical.frl) to identify novel potential compounds that share similar chemical features when aligned in 3D space. We also generated a pharmacophore model for the Rv2421c-DND-MG complex that spanned the DND active site contact region and included one hydrogen bond and two hydrophobic contacts, using the on-line tool ZINCPharmer [14], which was then used to screen the ZINC database for potential lead compounds. The Pharmer technology implemented in ZINCPharmer is a high-performance search engine that employs novel search methods such as geometric hashing, generalized Hough transforms and Bloom fingerprints to perform an exact pharmacophore match which accelerates the search algorithm [14]. The searches yielded 14 compounds after screening the PCL database, and 141 compounds after screening the ZINC database for purchasable compounds, which were further prioritized using docking studies.

**Fig 1:**
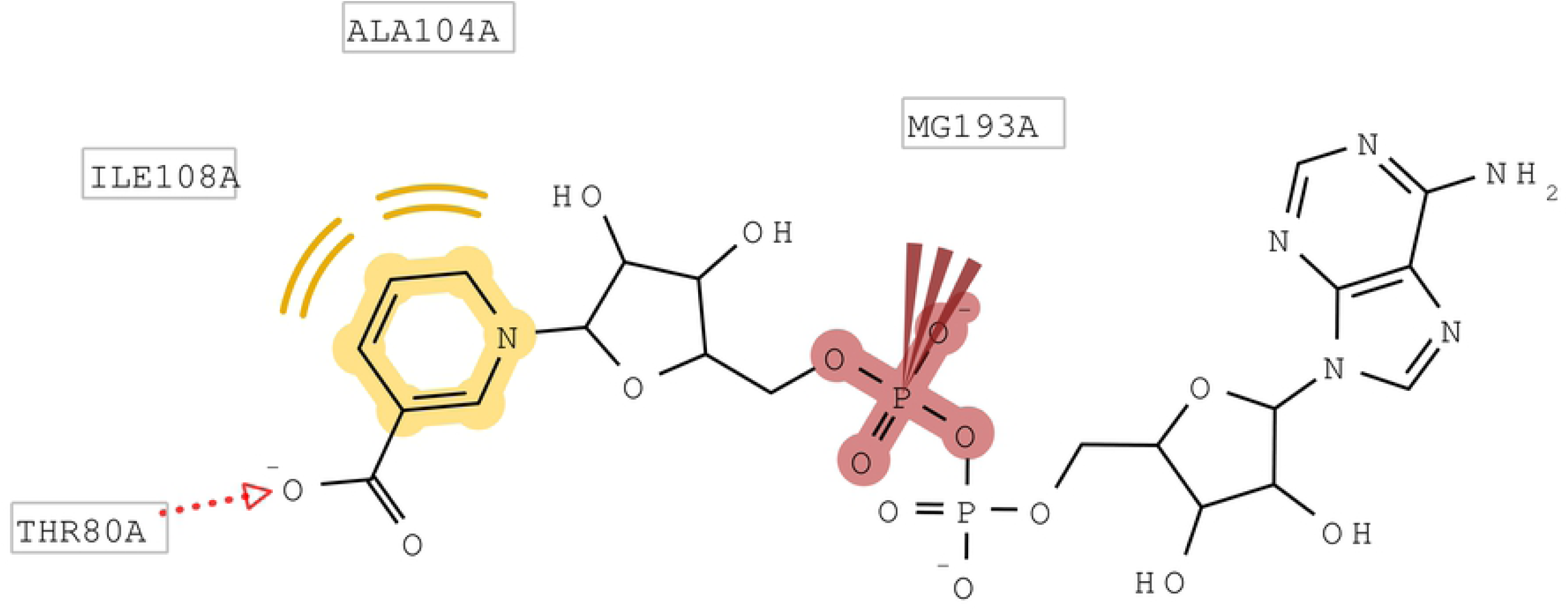
2D diagrams showing Pharmacophoric feature selection. A) model with four features and B) Model with two features for Rv2421c_DND_MG generated using LigandScout.

### 3-D Quantitative Structure Activity Relationship (QSAR) studies

The inhibitory efficiency of the obtained inhibitors was evaluated using 3D-QSAR studies. Known-TB NadD inhibitors were identified from the literature [15] to use as a training set to test our pharmacophore model. The structures of inhibitors were constructed and minimized using utilities of Discovery Studio (DS). The inhibitors were primarily processed using “Prepare Ligands for QSAR” module and the IC50 values were selected as the dependent variable. The whole dataset was further divided into the “Training” and “Test” sets and a 3D-QSAR model was constructed with 5-fold cross validations.

### Molecular docking using AutoDock Vina

155 compounds were identified with LIGANDSCOUT and ZINCPharmer, and these were docked to the energy minimized structure of Rv2421c using VINA [16]. The active site region spanning residues Thr8, Thr80, Thr100 and Leu157 was used for docking. A grid box size of (X= 26, Y =24 and Z=26) was sufficient to capture the whole active site region and an exhaustiveness search algorithm setting of 40 was used for the docking simulations. The grid setup is calculated automatically by VINA. VINA was able to reproduce the experimental binding pose as the top scoring pose of the ligand DND when benchmarked using the homologous template 3E27. As a comparative standard we also docked the known inhibitor ZINC58655383 of nicotinate mononucleotide adenylyltransferase (BaNadD for the *B*.*anthracis* (strain ∼CDC 684/NRRL 3496) to Rv2421c. The receptor and ligand DND were prepared using AutoDock tools which adds polar hydrogens, derives gasteiger charges for the atoms and saves the output file in pdbqt format. The 155 compounds and ZINC58655383 were prepared with automated bash and python scripts namely, split_multi_mol2_file.py and prepare_ligand4.py. After receptor and ligand preparation, the center of mass was calculated for the receptor structure with the ligand DND present. The various parameters for the docking process were stored in configuration files. The configuration file contained input parameters for the docking simulations such as the center of mass coordinates, grid dimensions and exhaustiveness of the search algorithm. The docked complexes were ranked according to their energy scores using the python script developed by the Scripps research institute vina_screen_get_top.py. The compounds that showed comparable and higher binding affinities than substrate DND and/or ZINC58655383 were visually inspected in PyMol and were further analyzed using PoseView [17]. PoseView determines four types of interactions namely i) hydrogen bonds, ii) hydrophobic interaction/bonds, iii) metal interactions and iv) π stacking interactions.

### Molecular dynamics simulations

The top scoring complexes were subjected to MD simulations using GROMACS 5.1.2 [18], with the topologies of the protein structures generated using the GROMOS96 53a6 force field [19]. The PRODRG server [20] was used to generate GROMOS96 based topologies as well as coordinate files of the inhibitors. The partial charges were corrected using the DFT method implemented in GAUSSIAN which utilized the B3LYP 6-31G (d,p) basis set and the CHELPG program [21]. Subsequently, all the docked complexes were solvated using the SPC/E water model [22] and the net charge in each box was neutralized by adding appropriate numbers of sodium (NA) and chloride (CL) counterions. The neutralized systems were energetically minimized by steepest descent and conjugate gradient algorithms.

Molecular dynamics simulations were carried out in the NVT (constant volume) and NPT (constant pressure) ensemble conditions, each for 100 ps during equilibration. The temperature of the system was maintained at 300 K using the Berendsen weak coupling method while the pressure was maintained at 1 bar by utilizing Parrinello-Rahman barostat. The production MD simulations were carried out for 100 ns. The generated trajectories were used to analyze the behavior of each complex for the last 50 ns of the simulation trajectory. The deviations in the distances, H-bonds, RMSD (Root Mean Square Deviations), and Radius of Gyration (Rg) were analyzed between the protein and ligands. The free energy of binding between the protein and ligands was calculated using the Molecular Mechanics Poisson–Boltzmann Surface Area (MM-PBSA) protocol implemented in the *g_mmpbsa* package for the last 50 ns of the trajectory [23].

### Experimental whole cell assays

#### Compound preparation

The top five compounds based on binding affinity scores and number of favorable interactions were purchased from chemical vendors (Sigma Aldrich, South Africa; Molport, USA; MCULE, USA, VITASMLAB, SPECS). Water insoluble compounds were dissolved in dimethyl sulfoxide (DMSO) (Merck Laboratories, USA) to obtain a final concentration of 800, 400, 200, 50, 25 and 10μM. Subsequently, 100 µl of each compound was added to the appropriate wells of a 96 well flat bottom microtiter plate.

#### Bacterial strains and growth conditions

*M. tuberculosis* H37Rv (ATCC 27294) with (H37Rv:mCHERRY) or without (H37Rv) the pCHERRY3 reporter plasmid [24] was cultured in Middlebrook 7H9 liquid media (Becton Dickinson, USA) supplemented with 0.2% (v/v) glycerol (Merck Laboratories, USA), 0.05% Tween 80 (Sigma-Aldrich, Germany) and 10% albumin-dextrose-catalase (ADC) (Becton Dickinson, USA), in filtered screw cap tissue culture flasks (Greiner Bio-one, Germany). Hygromycin B (50µg/ml) was included for plasmid maintenance, where required. The cultures were incubated at 37°C until an optical density (OD_600_ = 1.0) was reached.

#### *In vitro* bioluminescent reporter assay

Each culture was strained (40µM filter; Becton Dickinson, USA) and diluted till a final OD_600_ of 0.02 (100µl) was added to each of the wells containing a final compound concentration of 800, 400, 200, 50, 25 and 10μM. Positive, negative and compound controls were included on each plate. The positive control contained 100µg/ml rifampicin and H37Rv:pCHERRY3 reporter. The negative controls included wells containing media only to identify contamination and an untreated fluorescence control with DMSO to confirm that the DMSO used to dissolve the compounds did not affect growth. A compound control was included similar to the final concentration of each compound in the absence of *M. tuberculosis* to confirm no autofluorescence. The background control contained *M. tuberculosis* H37Rv (without reporter) which is used to subtract the fluorescence measurement to gain the inherent fluorescence of H37Rv. An undiluted H37Rv:pCHERRY3 control was used for the experiment. Plates were incubated at 37°C and readings were taken at 0hour and 24-hour intervals thereafter for approximately five days using the BMG Labtech POLARstar Omega plate reader (BMG Labtech, Germany, excitation: 587 nm, emission: 610 nm).

## Results and Discussion

### Sequence and structural analysis of Rv2421c (PDB: 4XOE)

Sequence-structural alignments between the target 4X0E and homologous protein structures NAMN adenylyltransferase from *B. anthracis* (3E27) and nicotinate mononucleotide adenylyltransferase from *B. anthracis* (2QTN) indicate approximately 40% sequence identity and 85% sequence coverage. Most importantly, 100% conservation of active site residues was observed between the proteins, suggesting that recognition of the ligand DND would be dominated by the same forces (S1 Fig). The structural similarity was less than 2 Å between 4X0E and 3E27 (∼1.3 Å), and between 4X0E and 2QTN (1.5 Å) (S2 A and S2 B Figs). The only major difference between the structures are an over-closed 3_10_ helix topology in *M. tuberculosis* Rv2421c which prevents ATP binding and NAMN binding, thus rendering it inactive [8]. The 211 amino acid structure of Rv2421c (4X0E) consists of 10 alpha helices, 5 beta sheets and 15 coils (Fig2).

**Fig 2:**
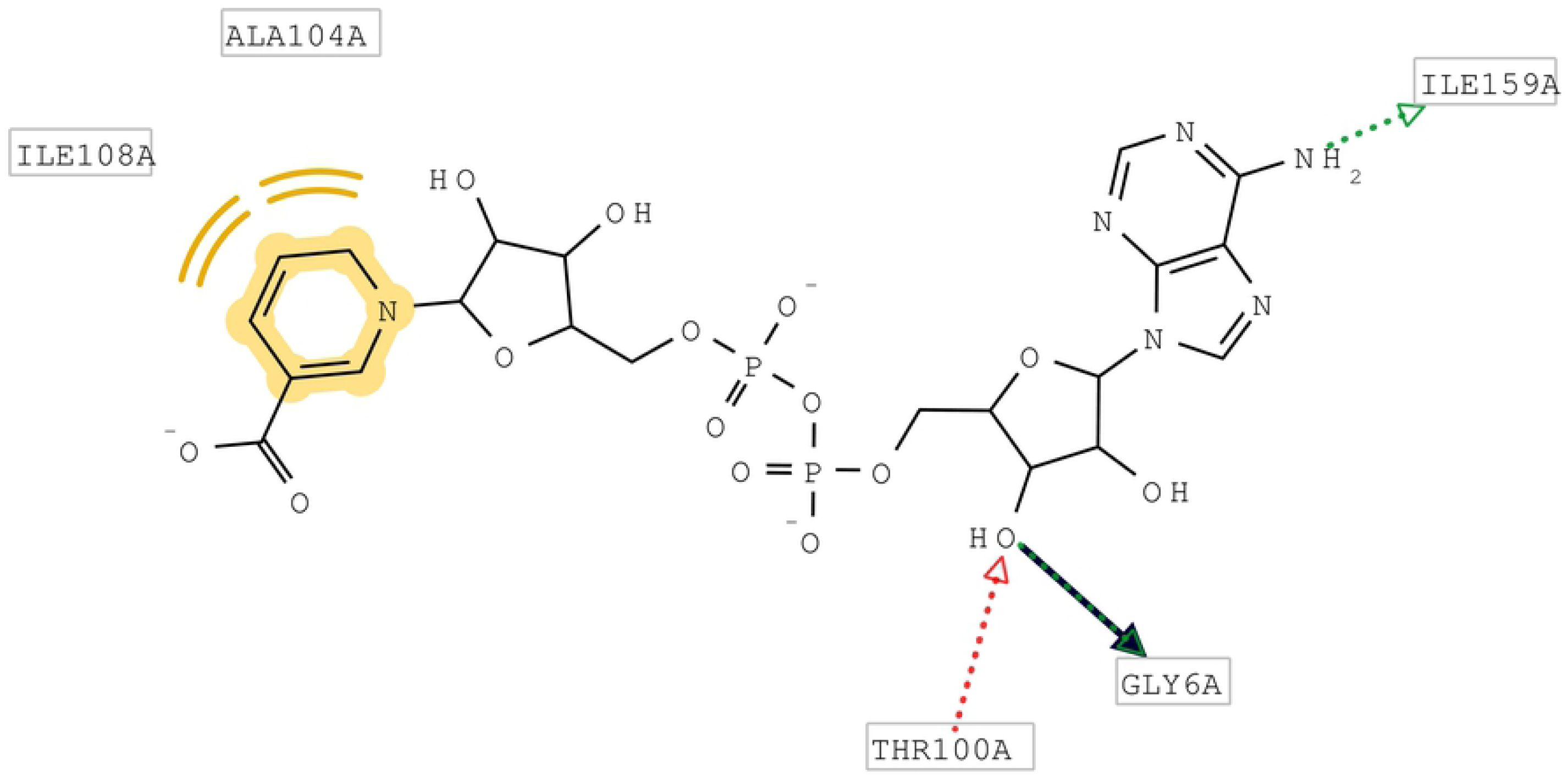
Cartoon representation of the three dimensional structure for Rv2421c in complex with DND and MG. The 3-10 over-closed helix conformation is shown in the box and the DND ligand is shown as sticks and MG as a sphere.

### 3D-QSAR analysis of the pharmacophore model

The non cross squared correlation coefficient (R^2^) and cross squared correlation coefficient (Q^2^) parameter values for the predicted 3D-QSAR model was calculated to be around 0.99 and 0.536 respectively, which was indicative of the reliability of the generated 3D-QSAR model. The activities of five molecules purchased for experimental testing; Carbenoxolone disodium salt (ZINC3977823), novel ZINC13544129, Cromolyn disodium salt (ZINC1530788), Novobiocin sodium salt (ZINC76945632) and Sulfasalazine (ZINC3831490) were computed to be around 7.92 µM, 5.97 µM, 6.93 µM, 5.81 and 4.99 µM respectively using the generated 3D-QSAR model. The outcome generated from this study indicated that these studied molecules have potential to inhibit TB Rv2421c protein functionality.

### Docking using VINA identifies three compounds that bind stronger than the known drug ZINC58655383 (LJZ)

The 155 compounds identified from the virtual screen against the Prestwick Chemical library and ZINC database were further filtered using VINA by docking to the energy minimized conformation of the protein. In total three compounds showed higher binding affinity values compared to the known drug ZINC58655383. These compounds included; novel ZINC13544129 and two FDA approved compounds used for the treatment of other conditions (Novobiocin sodium salt, Sulfasalazine, Table 1). The other two FDA approved compounds selected for this study, Carbenoxolone disodium salt and Cromolyn disodium salt (ZINC1530788), showed lower binding affinities compared to DND and ZINC58655383(Table 1).

**Table 1.** Docking scores and the number of interactions formed between the top 5 compounds, DND, known drug and binding site residues of Rv2421c.

| Structure | Number | Compound/<br>Substrate | Docking score<br>(Kcal/mol) | Number of HB | Number of<br>HPB | Number of<br>ionic contacts | Number of<br>π-π<br>interactions |
| --- | --- | --- | --- | --- | --- | --- | --- |
| Rv2421c | 1 | DND | -10.8 | 8 (Gly11, **Thr12**, Phe13, Asp14, **His20**, Asp77, **Ser168**, Thr169) | 1 (**Ser41**) | **1(MG)** | 0 |
|  | 2 | ZINC13544129 | -10.8 | 5 (**His20**, Thr86, Thr89, Gly107, Ser168) | 3 (**Ser41**, Gly107, Leu164) | 0 | 0 |
|  | 3 | Novobiocin disodium salt | -10.7 | 4 (**His20**, Pro83, Thr84, Gly107) | 2 (Ala110, Ile114) | **1(MG)** |  |
|  | 4 | Sulfasalazine | -9.6 | 4 (**Thr12**, Thr84, Thr86, **Ser168**) | 1 (**Ser41**) | 0 | 1 (Trp45) |
|  | 5 | ZINC58655383 | -9.5 | 0 | 3 (Trp45, Ala110, Leu164) | 0 | 1 (Trp45) |
|  | 6 | Carbenoxolone disodium salt | -9.1 | 2 (Thr45, Arg134) | 3 (**Thr12**, Arg45, Val51) | 0 | 0 |
|  | 7 | Cromolyn disodium salt | -8.6 | 3 (**Phe13**, **His20**, **Ser168**) | 2 (Ala110, Gly107) | 0 | 0 |

The number before curly bracket is total number of interactions. Abbreviations: DND-nicotinic acid adenine dinucleotide, HB- hydrogen bonds, HPB- hydrophobic bonds, Ala- Alanine, Arg- arginine, Asp- Aspartate, Gly- glycine, His- Histidine, Ile- Isoleucine, Leu- Leucine, MG- Magnesium, Phe- Phenylalanine, Pro- Proline, Ser- Serine, Thr- Threonine, Trp- Tryptophane, Val- Valine. Residues highlighted in bold are common interacting residues identified for each drug/substrate molecule.

### Interaction analysis of the five compounds selected for experimental validation

To further understand the mode of interaction between Rv2421c and DND, known inhibitor ZINC58655383 and the five compounds we performed interaction analysis using Poseview (S3 Figs A-G). The type of interactions is shown in Table 1 and in the Poseview two-dimensional interaction figures. The number of interactions formed between the novel ZINC13544129 and the four FDA approved compounds (Novobiocin sodium salt, Sulfasalazine, Carbenoxolone disodium salt and Cromolyn disodium salt) and residues of Rv2421c were 8, 7, 6, 5, and 5, respectively (S3 Figs E, F, D, B and C, Table 1).

### Analyses of conformational dynamics of Rv2421c and the five studied inhibitor docked complexes using MD simulations

Six 100 ns MD simulations were performed within a SPC/E immersed water model, minimized and equilibrated docked systems. The protein backbone stabilities of the systems were analyzed using RMSD analysis and these values showed equilibrium is reached after 50 ns in complex with inhibitors (S4 Figs A-F). The distance between the Rv2421c and each of the five compounds was calculated using gmx_mpi pairdist. The fluctuations in the distances were observed between 0.1 nm – 0.2 nm (Table 2). While for DND, Sulfasalazine and Novobiocin the computed distance values may reach below 0.16 nm (Table 2). Furthermore, the complex of DND showed an average number of 7.63 hydrogen bonds with Rv2421c active site residues and MG, followed by sulfasalazine and Cromolyn disodium up to 6.03 and 6.73 average number of hydrogen bonds formed between Rv2421c residues and MG, respectively (Table 2). The Novobiocin compound showed 4.25 hydrogen bonds in the system while Carbenoxolone and ZINC13544129 complexes contained less than three number of hydrogen bonds (Table 2). The compactness of the systems was assessed using the radius of gyration curves, which showed relatively similar range of compactness in the studied systems (S5 Figs A-E), with all converging towards 1.55 nm except for Carbenoxolone and Novobiocin systems each fluctuating around 1.6 nm.

**Table 2.** List of average values calculated for each of the MD simulation parameters for the last 50 ns.

| No | Compound name | Average values for last 50 ns |  |  |  |  |
| --- | --- | --- | --- | --- | --- | --- |
|  |  | Distance (nm) | Hydrogen Bonds | Total Energy (kJ/mol) | Van der Waals energy | Electrostatic Energy |
| 1. | DND | 0.169 ± 0.012 | 7.63 ± 2.02 | -185.52 ± 84.72 | -352.70 ± 31.27 | 167.18 ± 70.88 |
| 2. | Carbenoxolone | 0.174 ± 0.015 | 2.34 ± 0.86 | -29.75 ± 92.45 | -159.46 ± 40.20 | 129.71 ± 69.81 |
| 3. | Cromolyn disodium | 0.159 ± 0.010 | 6.73 ± 1.65 | -418.88 ± 91.88 | -182.82 ± 41.33 | -236.05 ± 68.21 |
| 4. | Sulfasalazine | 0.162 ± 0.014 | 6.03 ± 1.82 | -330.13 ± 53.10 | -237.45 ± 19.27 | -92.67 ± 51.22 |
| 5. | ZINC13544129 | 0.199 ± 0.022 | 2.13 ± 1.48 | -89.11 ± 69.83 | -233.69 ± 32.57 | 144.58 ± 50.72 |
| 6. | Novobiocin | 0.160 ± 0.013 | 4.25 ± 1.41 | -379.19 ± 26.54 | -318.62 ± 22.60 | -60.56 ± 14.09 |

The structure of Rv2421c contain an ATP active sub-site at His17, His20, Leu164, Ser167, Thr169, Arg172, while the residues Thr12, Asp14, Ser41, Ser52, Arg57, Thr86 form the NaMN active sub-site. The secondary structure corresponding to these active site residues are L1, A1, L3, A2, A4, A8, L13 and A9 (S6 Fig A). The fluctuations in the constituent residues were also studied which showed DND, Carbenoxolone and ZINC13544129 had more interactions with Rv2421c protein compared to Cromolyn disodium salt, Sulfasalazine and Novobiocin sodium salt (S6 Figs B-G). Furthermore, the interaction energies contribution to the binding between the Rv2421c and studied potential inhibitors were analyzed using MMPBSA protocol (Table 2). For Cromolyn disodium system, the highest total energy was calculated to be around −418.88 kJ/mol followed by Novobiocin and Sulfasalazine with the energy reaching up to −379.19 kJ/mol and −330.13 kJ/mol, while DND showed −185.52 kJ/mol (Table 3). Except, for the system where the major contributor to the total energy is Van der Waals interactions, and the rest being electrostatic energy (Table 2). These findings indicate the nature of binding between the Rv2421c and the studied complexes. The generated outcomes emphasized the need for further validation which are discussed in subsequent sections. In summary, Novobiocin showed closer distance to Rv2421c active site residues and higher binding free energy compared to known inhibitor ZINC13544129 and substrate DND making it a stronger competitive inhibitor of Rv2421c.

### *In vitro* growth assays verify Novobiocin sodium salt as a potential inhibitor of Rv2421c

We purchased five of the available top binding compounds (Carbenoxolone disodium salt, Novobiocin sodium salt, Sulfasalazine, Cromolyn sodium salt and ZINC13544129) based on higher binding affinity scores and higher number of interactions compared to DND and ZINC58655383. Unfortunately, no vendors were found for ZINC58655383 and DND, and these compounds were not purchased. In this study we aimed to identify inhibitors with MIC of at least 50 μM as a start for further interrogation. Of the five compounds (Fig3 A-E), only Novobiocin sodium salt was found to inhibit *M. tuberculosis* growth at 50, 25 μM and weakly at 10 μM concentrations (Fig 3B). Further inspection of the molecular properties and activity of Novobiocin sodium salt showed that it was slightly hydrophilic based on the milogP value of 0.66 and it had antibiotic activity against gram positive bacteria by inhibiting DNA synthesis.

**Fig 3:**
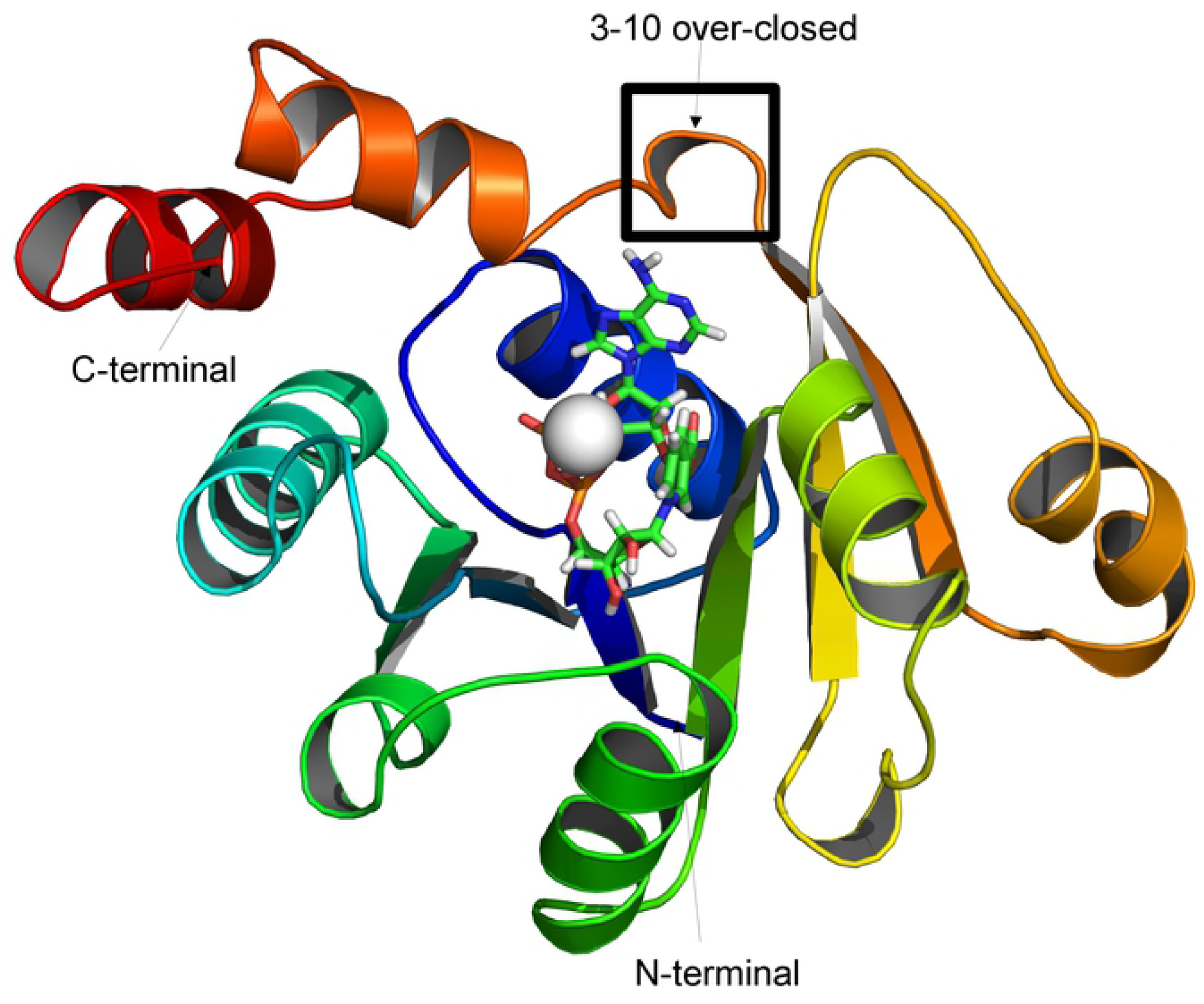
Survival growth curves for *Mycobacterium tuberculosis* at varying drug concentrations (10, 25, 50, 200, 400, 800 µM) for five days. Panel A = Carbenoxolone disodium salt, Panel B = Novobiocin sodium salt, Panel C = Sulfasalazine and Panel D = Cromolyn disodium salt, Panel E = ZINC13544129.

### Novobiocin a possible new mode of action for Rv2421c inhibition

The mechanism of action of Novobiocin includes the inhibition of DNA gyrase B and Topoisomerase IV in gram positive bacteria such as *Pseudomonas Aeruginosa, Acinetobacter baumannii* and *Klebsiella pneumonia* [25]. However, mutations in DNA gyrase B have resulted in resistance to Novobiocin specifically, in *Halophilic Archaebacteria* and *M. Tuberculosis* [26, 27]. Previous reports indicate that Novobiocin was withdrawn from the market due to poor pharmacological properties and safety concerns in the treatment of staphylococcus infections and therefore emphasized the need for alternative forms of the compound [25, 27]. Several derivatives of Novobiocin have been developed of which four were docked to Rv2421c. The docking results indicated that Chlorobiocin had the highest affinity (−11kcal/mol) followed by Coumermycin, compound 5 and BL-C43 (−10.4, −9.6 and −9.3 kcal/mol respectively). These derivatives could serve as alternatives for TB treatment. We constructed protein models for both Haloferax DNA gyrase B and *M. tuberculosis* DNA gyrase B using the Swissmodel webserver. Structural superimpositions indicated very low RMSD values (0.67Å) between the two models and the conservation of two mutant residues Ser122 and Arg137 suggesting that *M. tuberculosis* DNA gyrase B is resistant to Novobiocin. However, amino acid sequence comparisons between Haloferax DNA gyrase B and Rv2421c indicated very low, 10% sequence identity and only one conserved residue (Ser 122) associated with Novobiocin resistance in Haloferax. Additionally, Rv2421c is structurally different from *M. tuberculosis* DNA gyrase B with an RMSD value of more than 19Å. Molecular docking studies of Novobiocin and its four derivatives to *M. tuberculosis* DNA gyrase B indicated lower binding affinities for compound 5, Coumermycin, Novobiocin, Chlorobiocin and BL-C43 (−5.9, −6.5, −6.7, −6.7 and −6.8 kcal/mol respectively) compared to Rv2421c binding. This study provides data to suggest that Novobiocin derivatives can be considered for Rv2421c inhibition instead of the known gyrase B drug target because they bind to Rv2421c with higher affinity. Future work will include purchasing and or synthesizing the derivatives to test their activity against *M. tuberculosis* in whole cell assays, and possibly modifying the structure of Novobiocin sodium salt to improve MIC values.

## Acknowledgements

We would like to acknowledge the Centre for High Performance Computing (CHPC), Rondebosch, South Africa for allowing us to run our molecular dynamics simulations on their cluster. The authors also express their gratitude towards the Medical Research Council (MRC) and the National Research Foundation (NRF) for providing financial subsistence. RC and AC were funded by the South African Research Chairs Initiative of the Department of Science and Innovation (DSI) and the National Research Foundation (NRF) of South Africa (award number UID 64591). SLS was funded by the South African Research Chairs Initiative of the Department of Science and Innovation (DSI) and the National Research Foundation (NRF) of South Africa (award number UID 86539).

## Supporting Information

**S1 Fig. Multiple sequence alignment between Rv2421c (probable nicotinate-nucleotide adenylyl transferase (nadD), 212AA in size) and two homologous templates**.

PDBID: 2QTR crystal Structure of nicotinate mono-nucleotide adenylyltransferase from *B. anthracis*, PDBID: 3E27 Nicotinic acid mononucleotide (NaMN) adenylyltransferase from *B. anthracis*: product complex.

The alignment was performed using CLUSTALW v2.1 and was visually represented using Jalview v2.3. Shown in the red box are highly conserved signature sequence motif [(H/T)XGH]. DND active site residues are shown in black boxes.

**S2 Fig. Superimposition of the crystal structure of *M. tuberculosis* Rv2421c (4X0E) and homologous templates 3E27 and 2QTN**. A) The green structure represents Rv2421c and cyan is 3E27 solved in complex with nicotinic acid adenine dinucleotide (DND). Substrate DND (red) shown as a stick representation and MG (green) shown as a sphere RMSD = 1.261Å. B) The green structure represents Rv2421c and in pink is 2QTN solved in complex with nicotinate mononucleotide (NCN). Substrates DND (green) and NCN (cyan) shown as stick representations. RMSD = 1.502Å.

**S3 Fig. 2D interaction diagrams generated for the six complex systems using PoseView**. A) substrate DND is displaying eight hydrogen bond interactions, one hydrophobic contact and one MG ionic interaction with Rv2421c residues. B) Carbenoxolone disodium salt is displaying two hydrogen bond interactions and three hydrophobic contacts with Rv2421c residues. C) Cromolyn disodium salt is displaying three hydrogen bond interactions and two hydrophobic contacts with Rv2421c residues. D) Sulfasalazine is displaying four hydrogen bond interactions, three hydrophobic contacts and one pi-pi stacking interaction with Rv2421c active site residues. E) novel ZINC13544129 is displaying five hydrogen bond interactions and three hydrophobic contacts with Rv2421c residues. F) Novobiocin disodium salt is displaying four hydrogen bond interactions, two hydrophobic contacts and one MG ionic interaction with Rv2421c residues. G) The known NadD *E*.*coli* inhibitor ZINC58655383 is displaying three hydrophobic contacts and one pi-pi stacking interaction with the binding site residues of Rv2421c.The dashed lines represent hydrogen bonds and the green spline segments illustrates hydrophobic contacts and green dots represent pi-pi stacking interaction.

**S4 Fig. Root mean square deviation of backbone atoms for the six ligand complex systems over the last 50 ns**. A) DND, B) Carbenoxolone disodium salt, C) Cromolyn disodium salt, D) Sulfasalazine, E) ZINC13544129 and F) Novobiocin salt.

**S5 Fig. Radius of gyration of the backbone atoms for the six ligand complex systems over the last 50 ns**. A) DND, B) Carbenoxolone disodium salt, C) Cromolyn disodium salt, D) Sulfasalazine, E) ZINC13544129 and F) Novobiocin salt.

**S6 Fig. Root mean square fluctuation of the six complex ligand systems over over the last 50 ns for Rv2421c**. A) Structural depiction of active site residue fluctuation, B) DND, C) Carbenoxolone disodium salt, D) Cromolyn disodium salt, E) Sulfasalazine, F) ZINC13544129 and G) Novobiocin salt.

---

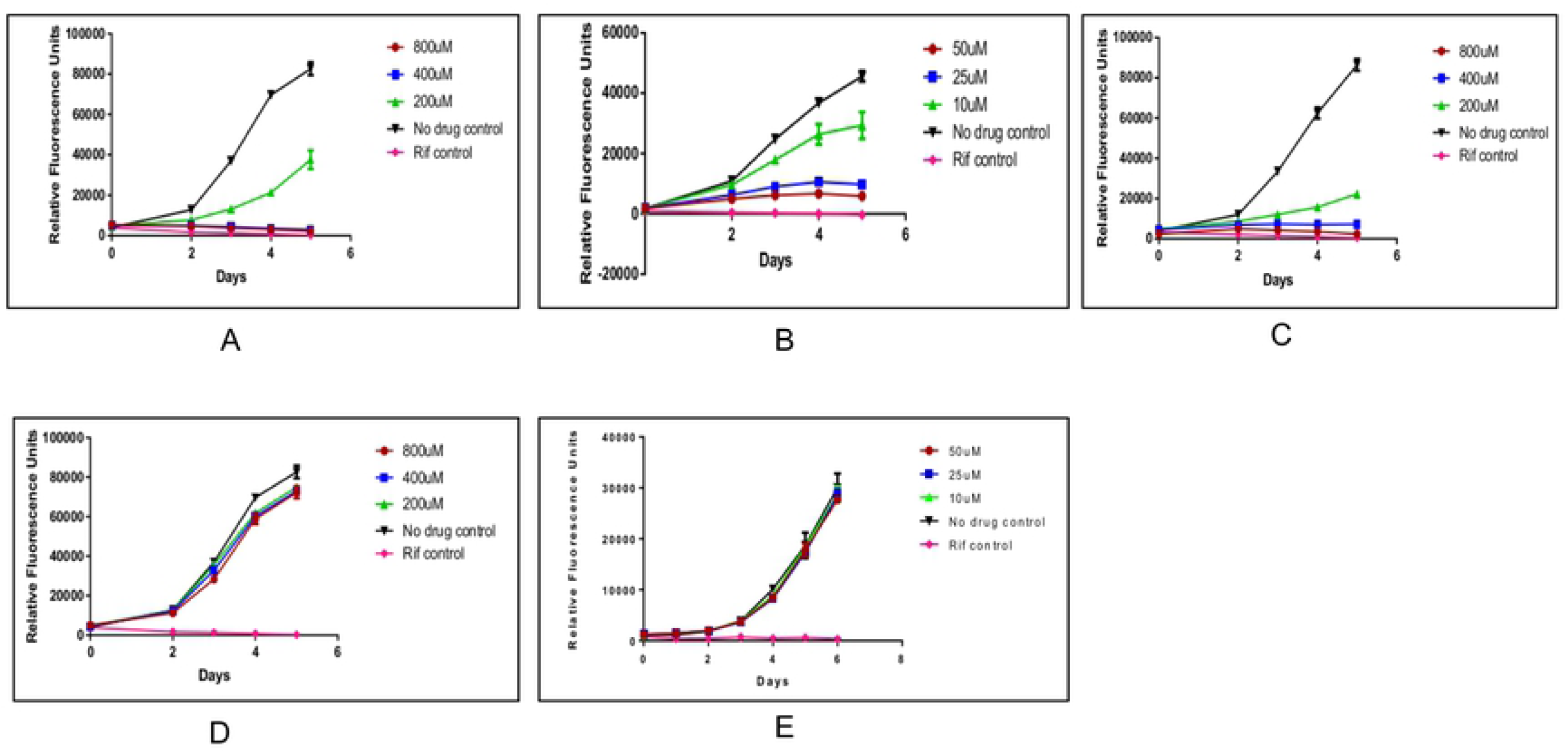

